# Acute Vaping of a golden Syrian Hamster is Feasible and Leads to Nicotine-Dependent Respiratory Tract Inflammation

**DOI:** 10.1101/2022.05.20.492852

**Authors:** Daniel M. Hinds, Heidi J. Nick, Tessa M. Vallin, Leslie A. Bloomquist, Sarah Christeson, Preston E. Bratcher, Emily H. Cooper, John T. Brinton, Angela Bosco-Lauth, Carl W. White

## Abstract

**Introduction:** E-cigarette vaping has become a major portion of nicotine consumption, especially for children and young adults. Although it is branded as a safer alternative to cigarette smoking, murine and rat models of sub-acute and chronic e-cigarette vaping exposure have shown many pro-inflammatory changes in the respiratory tract. An acute vaping exposure paradigm has not been demonstrated in the golden Syrian hamster, and the hamster is a readily available small animal model that has the unique benefit of becoming infected with and transmitting SARS-CoV-2 without genetic alteration to the animal or virus.

**Methods:** Using a two-day, whole-body vaping exposure protocol in male golden Syrian hamsters, we evaluated serum cotinine, bronchoalveolar lavage cells, lung and nasal histopathology, and gene expression in the nasopharynx and lung through RT-qPCR. Depending on the presence of nonnormality or outliers, statistical analysis was performed by ANOVA or Kruskal-Wallis tests. For tests that were statistically significant (p-value <0.05), post-hoc Tukey-Kramer and Dunn’s tests, respectively, were performed to make pairwise comparisons between groups.

**Results:** In nasal tissue, RT-qPCR analysis revealed nicotine-dependent increases in genes associated with type 1 inflammation (CCL-5 and CXCL-10), fibrosis (TGF-β), and a nicotine-independent decrease in the vasculogenesis/angiogenesis gene VEGF-A. In the lung, nicotine-dependent increases in the expression of genes involved in the renin-angiotensin pathway (ACE, ACE2), coagulation (tissue factor, Serpine-1), extracellular matrix remodeling (MMP-2, MMP-9), type 1 inflammation (IL-1β, TNF-α, and CXCL-10), fibrosis (TGF-β and Serpine-1), oxidative stress response (SOD-2), neutrophil extracellular traps release (ELANE), and vasculogenesis and angiogenesis (VEGF-A) were identified.

**Conclusion:** To our knowledge, this is the first demonstration that the Syrian hamster is a viable model of e-cig induced inhalational injury. In addition, this is the first report that e-cig vaping with nicotine can increase tissue factor gene expression in the lung. Our results show that even an acute exposure to e-cigarette vaping causes significant upregulation in the respiratory tract of pathways involving the renin-angiotensin system, coagulation, extracellular matrix remodeling, type 1 inflammation, fibrosis, oxidative stress response, NETosis, vasculogenesis, and angiogenesis.

## INTRODUCTION

E-cigarettes (e-cig), first introduced in on the market in 2003, have become a very popular form of nicotine consumption in the last several years. In 2021, 34% of high school students reported ever using tobacco products, and e-cig vaping represented 85% of their consumption (1). This means over 4.4 million high school students have tried e-cig vaping (1). Furthermore, just 40% of high school and middle school e-cig users report doing so on more than 20 of the last 30 days (1). This indicates that a significant portion of e-cig vaping is more intermittent and acute in duration. Marketed as a safer alternative to traditional cigarette use, the chemical content of the aerosol has significant overlap with the content of cigarette smoke as seen by the presence of small aldehydes like acrolein, acetaldehyde, and formaldehyde (2). These chemicals, along with the many others already identified in e-cig aerosol, play a part in the pro-inflammatory state created in the respiratory tract due to vaping. Evidence of the inflammation created by e-cig vaping has been shown in multiple murine and rat models. Especially in chronic and sub-acute exposures, e-cig exposure is known to cause lung lipid and macrophage dysregulation independent of nicotine, nicotine-dependent lung inflammation, extracellular matrix (ECM) remodeling, cytokine production, and alterations in response to reactive oxygen species (ROS) (3-5). Alterations in ROS response has been shown in-vivo and in-vitro in acute models, as well (6, 7). To our knowledge, golden Syrian hamsters (*Mesocricetus auratus*) have never been used as a model of e-cig induced inhalational injury. Since a substantial subset of e-cig vaping behavior is intermittent and acute, especially in teenagers, the development of models that reflect this type of exposure is important.

In this study, we hypothesized that the golden Syrian hamster would be a viable model of airway inflammation, ECM remodeling, oxidative stress, neutrophil extracellular traps release (NETosis), and risk of thrombosis related to acute e-cig exposure. Our results establish the validity of small animal modeling of e-cig inhalational injury in a hamster, and the novel finding that acute vaping leads to an increase in tissue factor expression. As an added benefit, the golden Syrian hamster is a readily available small animal model that does not require genetic modifications to become infected or transmit SARS-CoV-2 (8, 9). This may allow for the future study of the impact of e-cig vaping on COVID-19.

## METHODS

### Animals

The studies were approved by the Institutional Animal Care and Use Committee of the University of Colorado Anschutz Medical Campus. The investigators adhered to the National Institutes of Health Guide for the Care and Use of Laboratory Animals (1996) and research was carried out in compliance with the Animal Welfare Act. Six to eight-week-old male LVG golden Syrian hamsters (weight 93-140g; Charles River Laboratories, Wilmington, MA) were used in this study, maintained in an AAALAC-accredited animal care facility. Hamsters were acclimated for 7 days following arrival, housed four animals per cage, and their health status was monitored daily. Since this is a feasibility study of a novel exposure paradigm, no power calculation was performed, and sample sizes were constrained by the maximal number of animals that could be exposed at one time.

### Chemicals and e-cigarette liquid

With the goal of having consistent e-liquid content and aerosol generation, vape juice was made daily in the lab. Propylene glycol (PG), vegetable glycerin (VG), and nicotine free base (>99% GC) were purchased from Sigma-Aldrich and used as the components of the e-liquid. E-liquid consisting of 50% PG, 50% VG, with or without 25 mg/mL (2.5%) nicotine was prepared the morning of each exposure. This formulation is most compatible with the e-cig device and atomizer described below.

### Aerosol generation and animal exposures

E-cigarette aerosol was generated using an inExpose system (SCIREQ, Montreal, Quebec) equipped with a Joyetech eVic VTC Mini mod device, which provides consistent aerosol content and particle size though concurrent use of an atomizer consisting of a nickel-based, temperature-controlled coil with 0.15 Ohm resistance with stainless steel housing and cotton-based wick (sub-ohm coil, KangerTech, Shenzhen, China) (10). The puff profile used for the exposure included a total volume of 70 mL, an exposure time of 3.3 s, every 30 s (11). Hamsters were exposed to e-cigarette aerosol for 2 h per day for 2 consecutive days via a whole-body exposure system with 10 L chamber (SciReq). The e-cig aerosol generated was passed through the condensing chamber and pumped into the mixing chamber with a flowrate of 1.0 L/min. The vapor was diluted with air in the mixing chamber and then delivered into the whole-body exposure chamber where hamsters were separated by dividers. Simultaneously, the e-cig aerosol in the exposure chamber was exhausted by another pump with a flowrate of 3.0 L/min. Both the pumps were calibrated and adjusted before each exposure. Pumps were cleaned following each exposure to minimize the effects of nicotine residues. The atomizer was changed out after a maximum of 20 exposure hours to avoid overheating and carbon monoxide generation. Prior to exposure with a new atomizer, 1 mL of e-cig liquid was dispensed onto the atomizer ensuring saturation of the cotton wick and prevention of a dry burn. Hamsters were exposed in one of three groups: control animals (“bias flow” group), PG/VG + 0% nicotine, and PG/VG + 2.5% nicotine. Control animals received room air only, at a flowrate of 3.0 L/min. After 120 minutes of continuous, ambient exposure, animals were monitored for at least 15 minutes before returning to the housing facility.

### Euthanasia

Animals were euthanized immediately following the last day of e-cigarette exposure. Euthanasia was performed via terminal ketamine/ xylazine intraperitoneal (IP) injection (200 mg/kg ketamine + 30mg/kg xylazine; VetOne) followed by diaphragmatic puncture and exsanguination.

### Bronchoalveolar lavage fluid (BALF), lung tissue harvest and histopathology

After terminal anesthesia and exsanguination, lung tissue was cleared of blood by flushing the right ventricle with phosphate buffered saline (PBS) at 30 mL/min flow rate. The trachea was then cannulated, left bronchus clamped, and the right lung was lavaged with two 2.5 mL washes of PBS solution. Right lung lobes were tied off, individually dissected, immediately snap frozen in liquid nitrogen, and stored at -80°C until RNA extraction. Alternatively, the right upper lobe (RUL) was placed into 1 mL cold TRIzol Reagent (Invitrogen) per 100 mg of tissue, immediately homogenized, and the homogenate stored at -80°C until RNA extraction. The left lung was then unclamped and intratracheally fixed with 4% paraformaldehyde in PBS (PFA/ PBS) for 5 minutes prior to removal by gross dissection and storage in 4% PFA/PBS until embedding. Paraffin embedded tissues were sectioned at a thickness of 5 µm and stained for hematoxylin and eosin (H&E) and Alcian blue-periodic acid-Schiff (AB-PAS). BALF samples were pooled, centrifuged at 300 g for 10 min at 4 °C, and supernatants frozen (-80 °C). Cell pellets from BALF were re-suspended using 250 μL PBS and cytospin slides (Thermo Shandon) were prepared using 50,000 cells per slide. Differential cell counts (∼400 cells/slide) were performed on cytospin-prepared slides stained with PROTOCOL Hema 3 (Fisher Scientific). Total cell counts are expressed per microliter of BALF.

### Nasal respiratory epithelium tissue harvest and histopathology

After terminal anesthesia and exsanguination, nasal respiratory epithelial tissue was isolated by first removing the head at the base of the skull. The skin of the head and lower jaw was completely removed. Muscle and connective tissue were scraped from the frontal and nasal plates, the rhinarium was detached, and the nasal and frontal plates removed. The nasal respiratory epithelium and olfactory bulbs were collected and placed into 1 mL of cold TRIzol on ice. Samples were homogenized in TRIzol immediately following harvest and homogenates stored at -80°C until RNA extraction. For histopathology, the skin of the head and lower jaw was removed, along with brain matter through the back of the skull. The remaining skull was fixed in 150 mL of 4% PFA/PBS. After decalcification for 1.5 days, paraffin embedded tissues were sectioned at a thickness of 5 µm and stained for H&E and AB-PAS.

### Cotinine assay

Cotinine levels were measured in hamster serum collected immediately after the last day of e-cigarette aerosol exposure by ELISA according to the manufacturer’s instructions (Abnova). Approximately three ml of fresh blood was collected via inferior vena cava prior to exsanguination. The blood was allowed to clot, centrifuged at 1,300 g for 10 min at 25 °C, and the serum was collected, aliquoted, and stored at -80°C until analysis. Absorbance was measured at 450 nm with background correction at 560 nm using a SpectraMax M5 plate reader (Molecular Devices).

### Reverse transcription quantitative real-time PCR (RT-qPCR)

Total RNA was extracted using TRIzol Reagent and an RNeasy Mini Kit (Qiagen). After TRIzol-based isolation, RNA samples were equilibrated with an equal volume of 70% ethanol. A proportion of sample equivalent to ≤30 mg of tissue was then purified using the RNeasy Mini, including on-column DNase treatment, according to the manufacturer’s protocol. cDNA was synthesized from 50 ng of input RNA using the iScript Reverse Transcription Supermix (Bio-Rad), which contains a blend of oligo(dT) and random hexamers, according to the manufacturer’s instruction. Because commercial PCR panels were not available, primers were designed for gene targets of interest that could be internally validated, and primer sequences are shown in Table 1. The mRNA levels of our gene targets were detected using the SsoAdvanced Universal SYBR Green Supermix (Bio-Rad) on a QuantStudio 7 Flex system (Applied Biosystems, ThermoFisher). The manufacturer’s instructions for qPCR reaction preparation and thermal cycling parameters were followed. Target mRNA expression levels were normalized to Rpl18, which was validated for stable expression across all groups. Relative gene expression was determined by the 2^-ΔΔCt^ method and calculated relative to the mean expression level of the biological replicates of the bias flow control group. Data are presented as fold change compared to the bias flow control group. No reverse transcriptase controls and melt curve analyses were utilized to verify the absence of genomic amplification and specificity of amplicons.

**Table 1.**
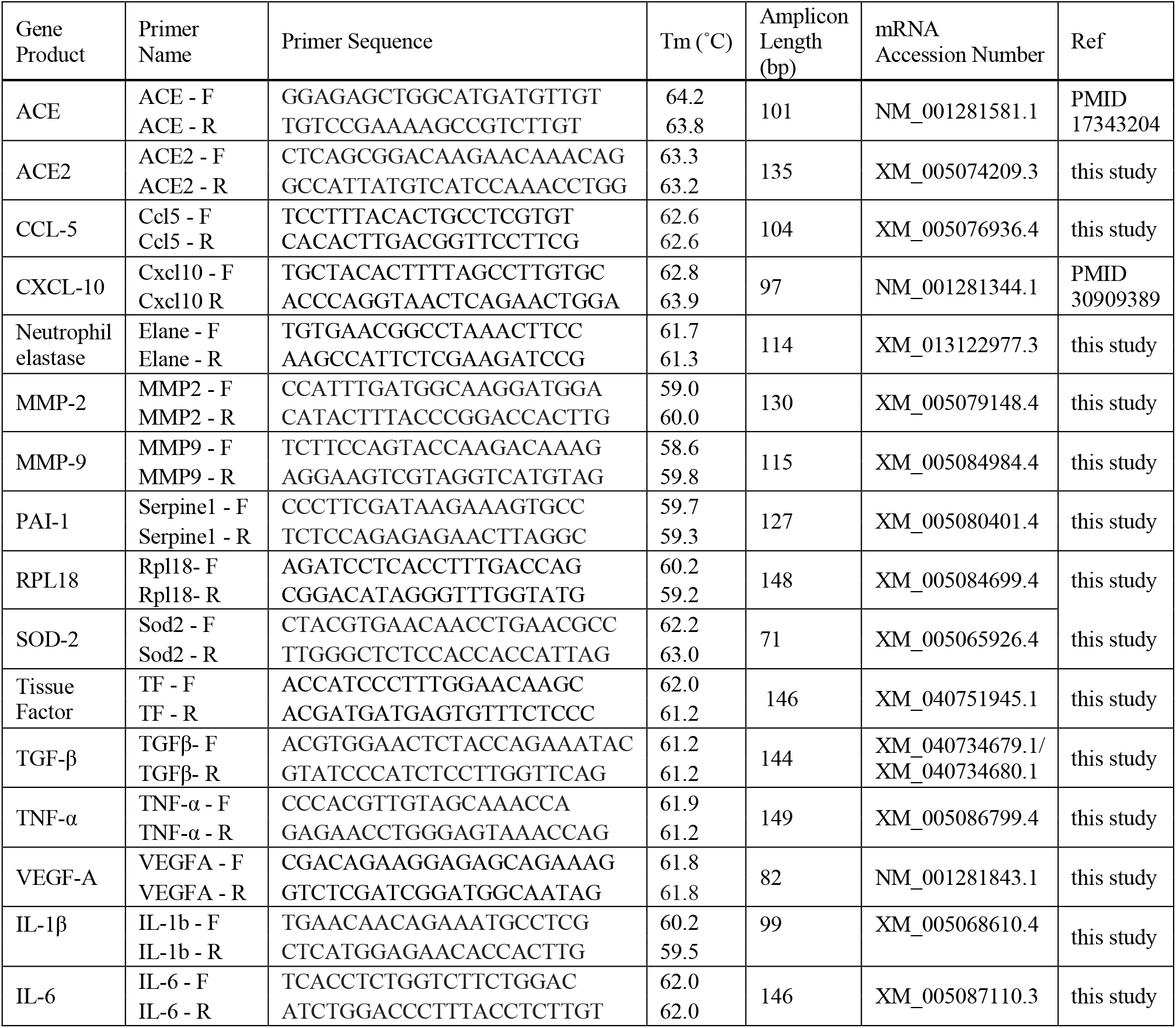
PCR primers.

### Data analysis and statistics

Based on exposure type (bias flow, PG/VG + 0% nicotine, and PG/VG + 2.5% nicotine), histograms and boxplots were used to visually identify outliers and check the distribution of outcome measures by group. There was no evidence of nonnormality or outliers in the BALF and cotinine data, thus ANOVA was used. Due to some outliers in our mRNA experiments of nasal respiratory epithelium and whole lung homogenate tissue, Kruskal-Wallis tests were used to compare fold change across groups. For the ANOVA and Kruskal-Wallis tests that were statistically significant (p-value <0.05), post-hoc Tukey-Kramer and Dunn’s tests, respectively, were performed to make pairwise comparisons between groups. To account for multiple comparisons, the familywise type I error rate for each set of 3 pairwise Tukey-Kramer tests was set to 0.05 and the p-values from each set of 3 pairwise post-hoc Dunn’s tests were adjusted using the Bonferroni correction. All data visualizations and analyses were performed with GraphPad Prism software (Version 9.2, La Jolla, CA).

## RESULTS

### Hamster behavior and clinical monitoring does not change with acute vaping

Hamster behavior and clinical monitoring was performed daily, at least 15 minutes after completion of their exposure. Using a multi-system checklist, we found no abnormalities, thus, no difference in behavior or clinical findings between the bias flow, PG/VG + 0% nicotine, and PG/VG + 2.5% nicotine groups were present.

### Cotinine is detectable in hamster serum after acute nicotine vaping

After two-days of exposure, both bias flow (n=7) and PG/VG + 0% nicotine (n=8) hamsters had plasma cotinine levels less than 5 ng/mL (Fig. 1A; 0.0 ± 0.0 ng/mL and 1.4 ± 0.7 ng/mL, respectively). Serum cotinine levels were significantly higher in hamsters exposed to PG/VG + 2.5% nicotine (n=7) (Fig. 1A; 62.3 ± 10.4 ng/mL; p < 0.0001).

**Legend – Figure 1 -.**
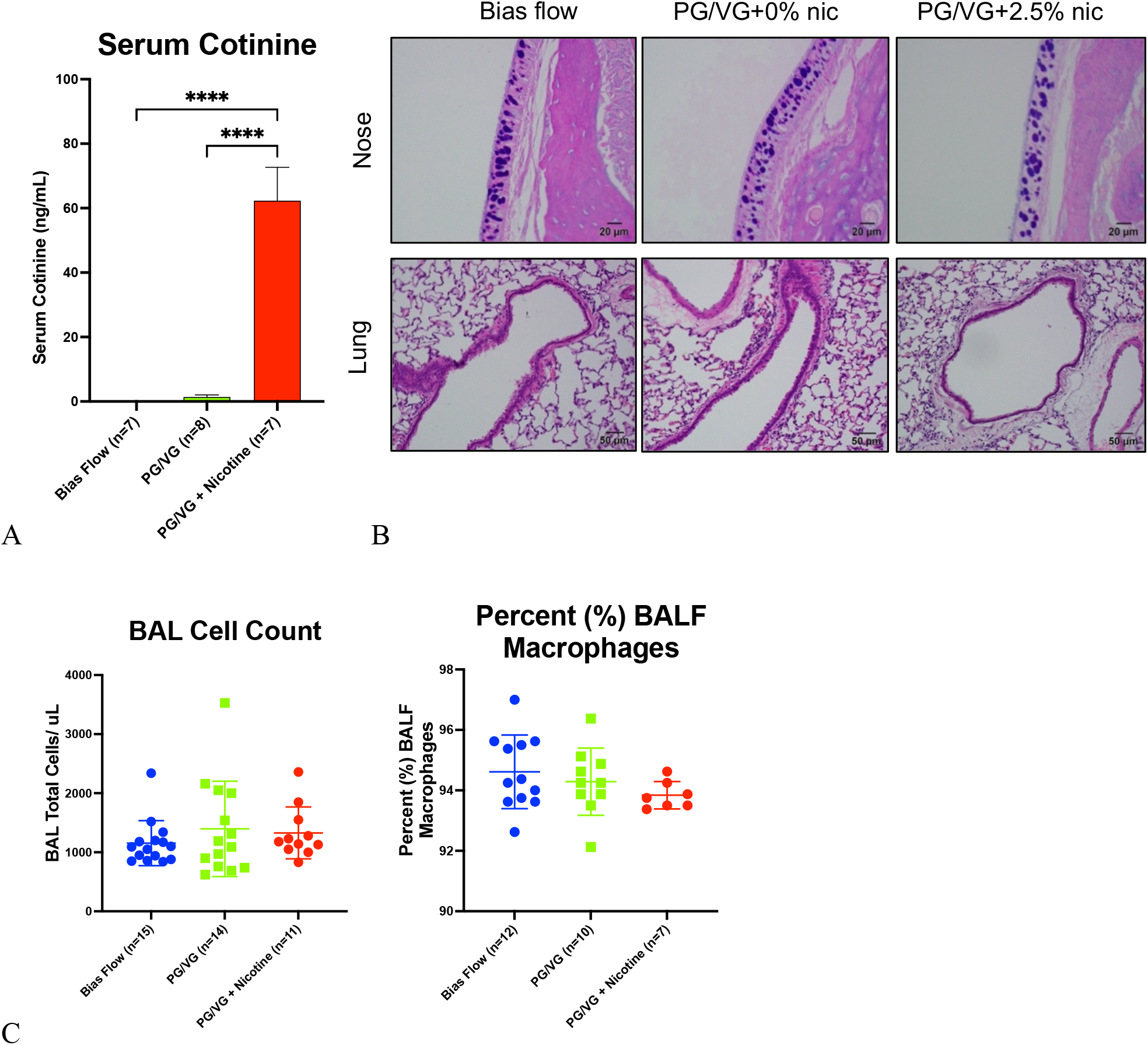
Acute, whole-body e-cigarette exposure. a) serum cotinine levels for animals exposed to bias flow (0.0 ± 0.0 ng/mL), PG/VG + 0% nicotine (1.4 ± 0.7 ng/mL), or PG/VG + 2.5% nicotine (62.3 ± 10.4 ng/mL). **** p < 0.0001, ANOVA with Tukey-Kramer post-hoc test. b)AB-PAS-stained nasal sections and H&E-stained lung sections from animals exposed to bias flow (left panels), PG/VG + 0% nicotine (middle panels), or PG/VG + 2.5% nicotine (right panels). c)total cell count and proportion of macrophages in bronchoalveolar lavage (BAL) fluid for animals exposed to bias flow (1154 ± 381 cells/uL), PG/VG + 0% nicotine (1396 ± 806 cells/uL), and PG/VG + 2.5% nicotine (1327 ± 439). BAL cell count p = 0.5171, percent BALF macrophages p = 0.3151, ANOVA with Tukey-Kramer post-hoc test. All data are reported as mean ± SD.

### Acute vaping does not cause histologically evident changes in mucus cell or epithelial organization in the hamster nose or lung

Histopathology of the nose and lung for each exposure group was evaluated after two days of exposure. In the nose, AB-PAS staining showed no discernable difference in mucin component production between the vaping regimens (Fig. 1B – AB-PAS Nose – bias flow, PG/VG + 0% nicotine, PG/VG + 2.5% nicotine). In the bronchial epithelium, H&E staining highlights that there were no significant differences in bronchial epithelial organization, sloughing, or inflammatory cell infiltration (Fig. 1B – H&E Lung – bias flow, PG/VG + 0% nicotine, PG/VG + 2.5% nicotine). Additionally, there were no differences in lung structure, architecture, or cell populations between the bias-flow group and the vaped groups (PG/VG + 0% nicotine or PG/VG + 2.5% nicotine). H&E staining of the nose and AB-PAS staining of the lung show similar findings to their respective counterparts (Supplemental Fig. 2).

### BAL cell numbers and composition were not altered by acute vaping

The total white blood cell (WBC) counts in BALF were not different between bias flow (n=15; 1154 ± 381 cells/μL), PG/VG + 0% nicotine (n=14; 1396 ± 806 cells/μL), and PG/VG + 2.5% nicotine (n=11; 1327 ± 439 cells/μL) (Fig. 1C, p = 0.52) after two days of vaping. Additionally, there was no difference in BALF cell differential as evidenced by the percent of macrophages in BALF between bias flow (n=12; 94.6 ± 1.2%), PG/VG + 0% nicotine (n=10; 94.3 ± 1.1%), and PG/VG + 2.5% nicotine (n=7; 93.8 ± 0.5%) (Fig. 1C, p = 0.32).

### Acute vaping alters nasal epithelial expression levels of genes regulating type 1 inflammation, fibrosis, and reactive oxygen species processing

Analysis of the impact of vaping on gene expression of the hamster airway was first evaluated by investigating changes in mRNA expression in nasal respiratory epithelium. We selected a panel of target genes (Table 1) based on relevance to SARS-CoV-2 infection/COVID-19 disease, as well as previously published reports of vaping-induced changes in the lungs of mouse models. For most genes investigated, there was no difference in expression level between the bias flow and PG/VG + 0% groups (Table 2, Fig. 2, and Supplemental Fig. 2). However, we found a nicotine-dependent increase in two genes representing a type 1 inflammatory response, CCL-5 (mean fold change + SD = 4.2 ± 3.3; p = 0.0101) and CXCL-10 (3.2 ± 1.5; p = 0.0009) (Table 2, Fig. 2A). Additionally, a nicotine-dependent increase in the pro-fibrotic mediator TGF-β was observed (1.6 ± 0.6; p = 0.0003; Table 2, Fig. 2B). The ROS response gene SOD-2 was unique in showing a nicotine-independent increase (PG/VG + 0% nicotine relative to bias flow 1.8 ± 0.5; p = 0.0133; Table 2, Fig. 2C), and a further increase with exposure to nicotine, although this was not statistically significant (PG/VG + 2.5% nicotine compared to PG/VG + 0% nicotine 1.8 ± 0.7; p = 0.1066; Table 2, Fig. 2C). This suggests that there are both PG/VG-related and nicotine-related effects on SOD-2 expression in the nasal epithelium. In contrast, we found a nicotine-independent decrease in VEGF-A, which regulates vasculogenesis and angiogenesis (PG/VG + 0% nicotine: bias flow = 0.5 ± 0.2; p = 0.0252 and PG/VG + 2.5% nicotine: bias flow = 0.4 ± 0.3; p = 0.0016; Table 2, Fig. 2D). For the type 1 inflammatory response gene TNF-α, expression was higher for the nicotine-exposed group compared to PG/VG alone, suggesting a nicotine-related effect, but this was not statistically significant (PG/VG + 2.5% nicotine: PG/VG + 0% nicotine = 1.7 ± 0.9; p = 0.1511; Table 2, Fig. 2A). Additionally, the mean expression of the type 1 inflammatory response gene IL-1β was slightly elevated for the PG/VG + 0% nicotine group compared to bias flow-exposed animals (1.4 ± 0.7; p = 0.9693; Table 2, Fig. 2A), a trend that was also observed in the comparison between the PG/VG alone and PG/VG + 2.5% nicotine groups (1.5 ± 0.8; p = 0.1887; Table 2, Fig. 2A). There was a significant increase in IL-1β expression between PG/VG + 2.5% nicotine and bias flow exposure (2.0 ± 1.1; p = 0.0132; Table 2, Fig. 2A). All other gene targets investigated showed no significant changes with the vaping of PG/VG + 0% nicotine or PG/VG + 2.5% nicotine (Table 2, Supplemental Fig. 2).

**Table 2.**
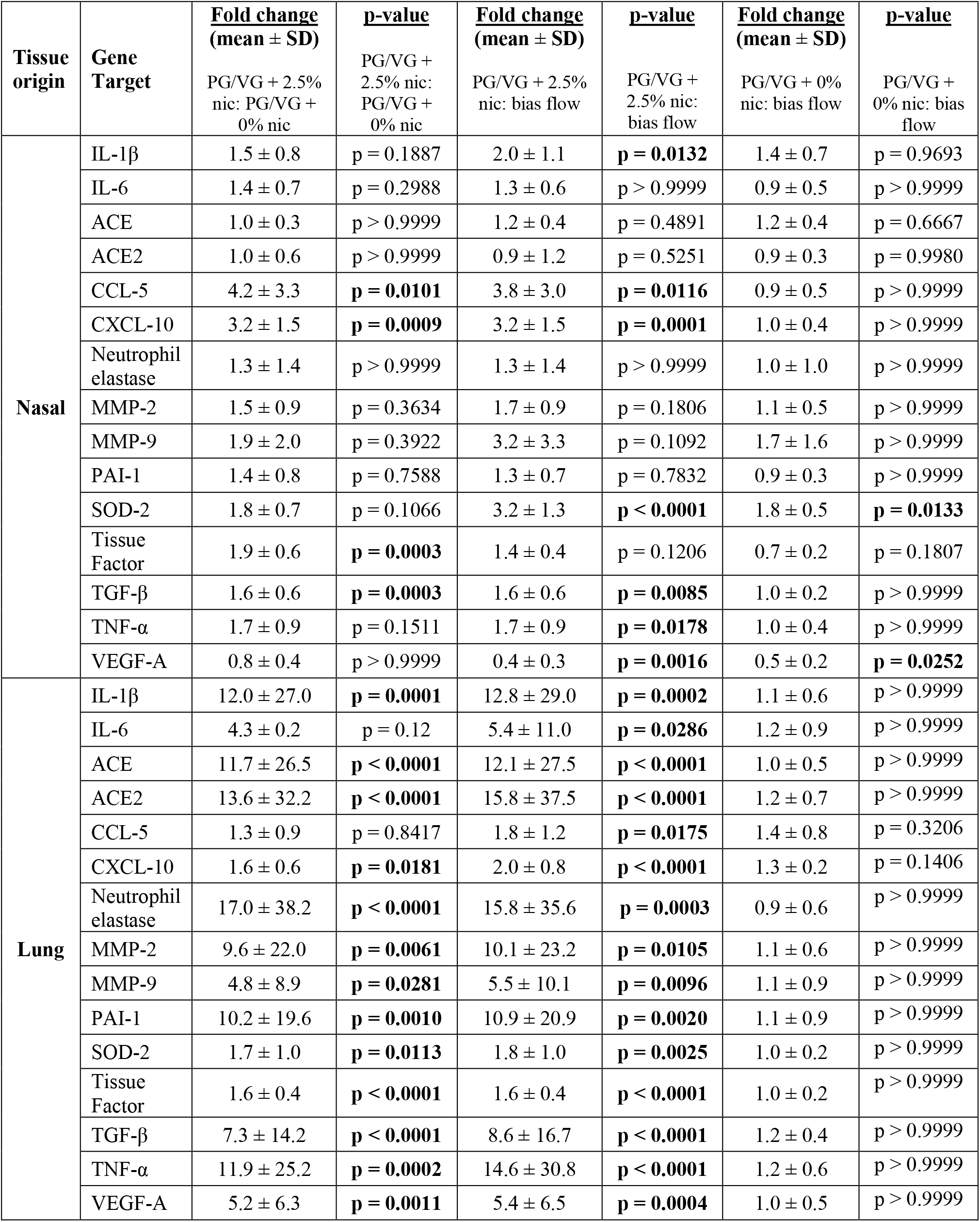
Respiratory Tract Gene Expression Changes with Acute Vaping Exposure.

| Tissue origin | Gene Target | <u>Fold change (mean ± SD)</u> | <u>p-value</u> | <u>Fold change (mean ± SD)</u> | <u>p-value</u> | <u>Fold change (mean ± SD)</u> | <u>p-value</u> |
| --- | --- | --- | --- | --- | --- | --- | --- |
|  |  | PG/VG + 2.5% nic: PG/VG + 0% nic | PG/VG + 2.5% nic: PG/VG + 0% nic | PG/VG + 2.5% nic: bias flow | PG/VG + 2.5% nic: bias flow | PG/VG + 0% nic: bias flow | PG/VG + 0% nic: bias flow |
| Nasal | IL-1β | 1.5 ± 0.8 | p = 0.1887 | 2.0 ± 1.1 | **p = 0.0132** | 1.4 ± 0.7 | p = 0.9693 |
|  | IL-6 | 1.4 ± 0.7 | p = 0.2988 | 1.3 ± 0.6 | p > 0.9999 | 0.9 ± 0.5 | p > 0.9999 |
|  | ACE | 1.0 ± 0.3 | p > 0.9999 | 1.2 ± 0.4 | p = 0.4891 | 1.2 ± 0.4 | p = 0.6667 |
|  | ACE2 | 1.0 ± 0.6 | p > 0.9999 | 0.9 ± 1.2 | p = 0.5251 | 0.9 ± 0.3 | p = 0.9980 |
|  | CCL-5 | 4.2 ± 3.3 | **p = 0.0101** | 3.8 ± 3.0 | **p = 0.0116** | 0.9 ± 0.5 | p > 0.9999 |
|  | CXCL-10 | 3.2 ± 1.5 | **p = 0.0009** | 3.2 ± 1.5 | **p = 0.0001** | 1.0 ± 0.4 | p > 0.9999 |
|  | Neutrophil elastase | 1.3 ± 1.4 | p > 0.9999 | 1.3 ± 1.4 | p > 0.9999 | 1.0 ± 1.0 | p > 0.9999 |
|  | MMP-2 | 1.5 ± 0.9 | p = 0.3634 | 1.7 ± 0.9 | p = 0.1806 | 1.1 ± 0.5 | p > 0.9999 |
|  | MMP-9 | 1.9 ± 2.0 | p = 0.3922 | 3.2 ± 3.3 | p = 0.1092 | 1.7 ± 1.6 | p > 0.9999 |
|  | PAI-1 | 1.4 ± 0.8 | p = 0.7588 | 1.3 ± 0.7 | p = 0.7832 | 0.9 ± 0.3 | p > 0.9999 |
|  | SOD-2 | 1.8 ± 0.7 | p = 0.1066 | 3.2 ± 1.3 | **p < 0.0001** | 1.8 ± 0.5 | **p = 0.0133** |
|  | Tissue Factor | 1.9 ± 0.6 | **p = 0.0003** | 1.4 ± 0.4 | p = 0.1206 | 0.7 ± 0.2 | p = 0.1807 |
| | TGF-β | 1.6 ± 0.6 | **p = 0.0003** | 1.6 ± 0.6 | **p = 0.0085** | 1.0 ± 0.2 | p > 0.9999 |
| | TNF-α | 1.7 ± 0.9 | p = 0.1511 | 1.7 ± 0.9 | **p = 0.0178** | 1.0 ± 0.4 | p > 0.9999 |
|  | VEGF-A | 0.8 ± 0.4 | p > 0.9999 | 0.4 ± 0.3 | **p = 0.0016** | 0.5 ± 0.2 | **p = 0.0252** |
| Lung | IL-1β | 12.0 ± 27.0 | **p = 0.0001** | 12.8 ± 29.0 | **p = 0.0002** | 1.1 ± 0.6 | p > 0.9999 |
|  | IL-6 | 4.3 ± 0.2 | p = 0.12 | 5.4 ± 11.0 | **p = 0.0286** | 1.2 ± 0.9 | p > 0.9999 |
|  | ACE | 11.7 ± 26.5 | **p < 0.0001** | 12.1 ± 27.5 | **p < 0.0001** | 1.0 ± 0.5 | p > 0.9999 |
|  | ACE2 | 13.6 ± 32.2 | **p < 0.0001** | 15.8 ± 37.5 | **p < 0.0001** | 1.2 ± 0.7 | p > 0.9999 |
|  | CCL-5 | 1.3 ± 0.9 | p = 0.8417 | 1.8 ± 1.2 | **p = 0.0175** | 1.4 ± 0.8 | p = 0.3206 |
|  | CXCL-10 | 1.6 ± 0.6 | **p = 0.0181** | 2.0 ± 0.8 | **p < 0.0001** | 1.3 ± 0.2 | p = 0.1406 |
|  | Neutrophil elastase | 17.0 ± 38.2 | **p < 0.0001** | 15.8 ± 35.6 | **p = 0.0003** | 0.9 ± 0.6 | p > 0.9999 |
|  | MMP-2 | 9.6 ± 22.0 | **p = 0.0061** | 10.1 ± 23.2 | **p = 0.0105** | 1.1 ± 0.6 | p > 0.9999 |
|  | MMP-9 | 4.8 ± 8.9 | **p = 0.0281** | 5.5 ± 10.1 | **p = 0.0096** | 1.1 ± 0.9 | p > 0.9999 |
|  | PAI-1 | 10.2 ± 19.6 | **p = 0.0010** | 10.9 ± 20.9 | **p = 0.0020** | 1.1 ± 0.9 | p > 0.9999 |
|  | SOD-2 | 1.7 ± 1.0 | **p = 0.0113** | 1.8 ± 1.0 | **p = 0.0025** | 1.0 ± 0.2 | p > 0.9999 |
|  | Tissue Factor | 1.6 ± 0.4 | **p < 0.0001** | 1.6 ± 0.4 | **p < 0.0001** | 1.0 ± 0.2 | p > 0.9999 |
| | TGF-β | 7.3 ± 14.2 | **p < 0.0001** | 8.6 ± 16.7 | **p < 0.0001** | 1.2 ± 0.4 | p > 0.9999 |
| | TNF-α | 11.9 ± 25.2 | **p = 0.0002** | 14.6 ± 30.8 | **p < 0.0001** | 1.2 ± 0.6 | p > 0.9999 |
|  | VEGF-A | 5.2 ± 6.3 | **p = 0.0011** | 5.4 ± 6.5 | **p = 0.0004** | 1.0 ± 0.5 | p > 0.9999 |

**Legend – Figure 2.**
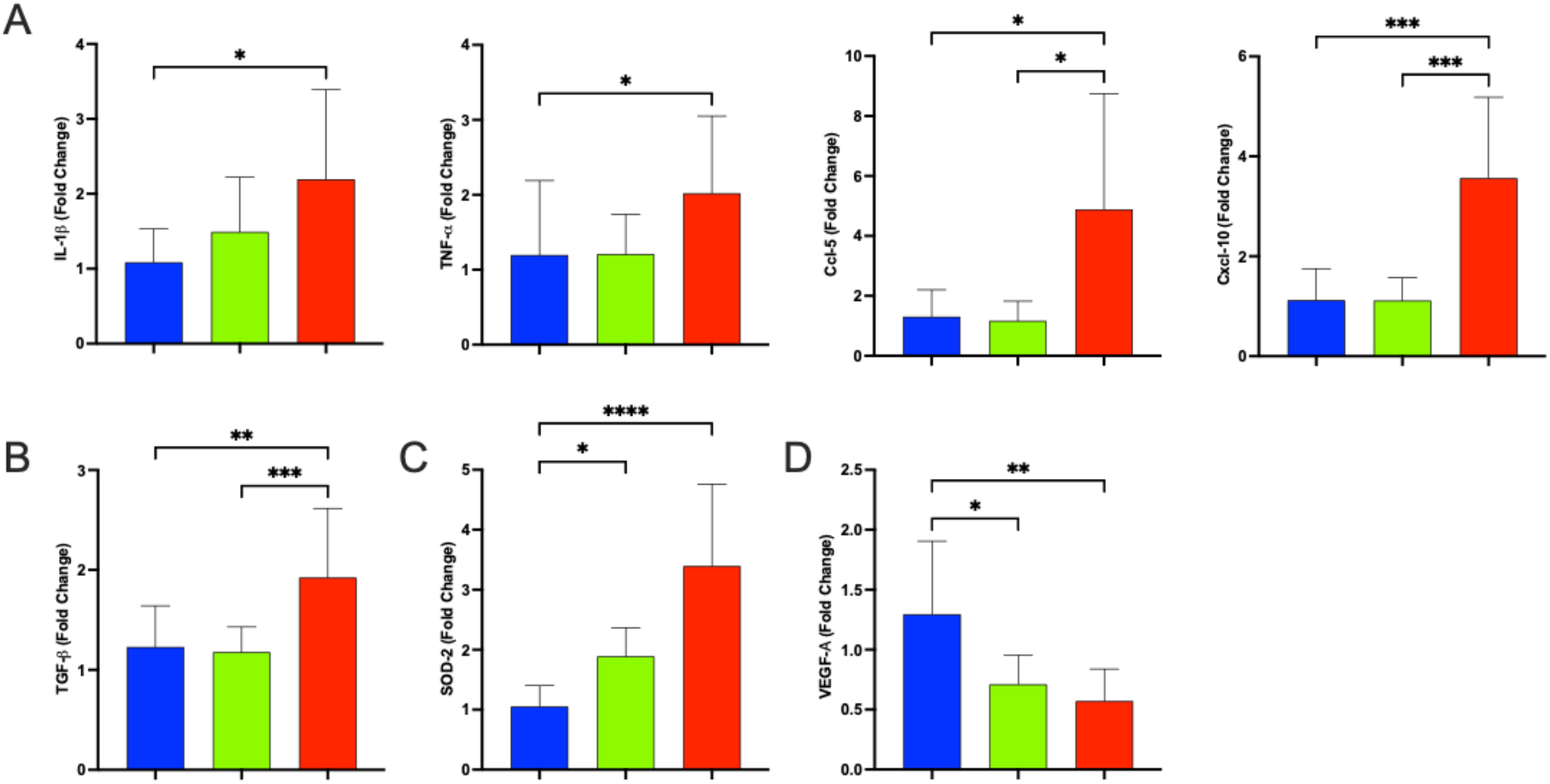
Changes in gene expression in nasal epithelia and olfactory bulb tissue induced by acute e-cig exposure. Hamsters were exposed to bias flow (blue), PG/VG + 0% nicotine (green), or PG/VG + 2.5% nicotine (red) for two days (n = 11-12/ group). Target gene expression was analyzed by RT-qPCR. Levels were normalized to Rpl18 and calculated relative to the mean expression level of the biological replicates of the bias flow group. (a) IL-1β, TNF-β, CCL-5, and CXCL-10, (b) TGF-β, (c) SOD-2, and (d) VEGF-A. *p<0.05, **p<0.01, ***p<0.001, **** p<0.0001, Kruskal-Wallis with Dunn’s post-hoc test.

### Vaping acutely alters lung expression of genes regulating inflammation, reactive oxygen species processing, coagulation, fibrosis, and repair response in a nicotine-dependent manner

To further characterize the impact of acute vaping on hamster airways, gene expression changes were evaluated in whole-lung homogenates. Thirteen of the 15 target genes investigated demonstrated a nicotine-dependent increase (Table 2, Fig 3A-G). This includes ACE (PG/VG + 2.5% nicotine: PG/VG + 0% nicotine 11.7 ± 26.5; p < 0.0001) and ACE2 (13.6 ± 32.2; p < 0.0001), genes involved in the renin-angiotensin (RAAS) pathway (Table 2, Fig. 3A). Notably, ACE2 also serves as the receptor for SARS-CoV-2. Exposure to nicotine also impacted the coagulation-related genes tissue factor (1.6 ± 0.4; p < 0.0001) and Serpine-1 (encoding PAI-1, 10.2 ± 19.6; p = 0.0010), extracellular matrix remodeling genes MMP-2 (9.6 ± 22.0; p = 0.0061) and MMP-9 (4.8 ± 8.9; p = 0.0281), and fibrosis and vasculogenesis genes TGF-β and VEGF-A (7.3 ± 14.2; p < 0.0001 and 5.2 ± 6.3; p = 0.0011, respectively) (Table 2, Fig. 3 B - D). Similar to the results obtained in the analysis of nasal epithelial tissue, several type 1 inflammatory genes were upregulated by nicotine vaping. These included IL-1β (12.0 ± 27.0, p = 0.0001), TNF-α (11.9 ± 25.2, p = 0.0002), and CXCL-10 (1.6 ± 0.6, p = 0.0181) (Table 2, Fig. 3E). In contrast to the pattern observed in nasal tissue, the ROS response gene SOD-2 was increased in a completely nicotine-dependent manner in lung tissue (1.7 ± 1.0; p = 0.0113; Table 2, Fig. 3F). Finally, a nicotine-dependent link to NETosis was evidenced by an increase in ELANE, the gene encoding neutrophil elastase (17.0 ± 38.2; p < 0.0001; Table 2, Fig. 3G). While there was no difference in IL-6 expression between the PG/VG alone and bias flow groups or between PG/VG alone and PG/VG + 2.5% nicotine, IL-6 was elevated in the nicotine-exposed group compared to bias flow animals (5.4 ± 11.0; p = 0.0286; Table 2, Fig. 3E). The mean expression of CCL5 was marginally elevated in the PG/VG alone compared to bias flow group (1.4 ± 0.8; p = 0.3206; Table 2, Fig. 3E), but there was an increase in its expression between PG/VG + 2.5% nicotine and bias flow exposure (1.8 ± 1.2, p = 0.0175; Table 2, Fig. 3E). CCL5 was not significantly different in the PG/VG alone versus PG/VG + 2.5% nicotine groups. None of the genes showed a significant change between PG/VG + 0% nicotine vaping and bias flow exposure (Table 2, Fig. 3A-E).

**Legend – Figure 3 -.**
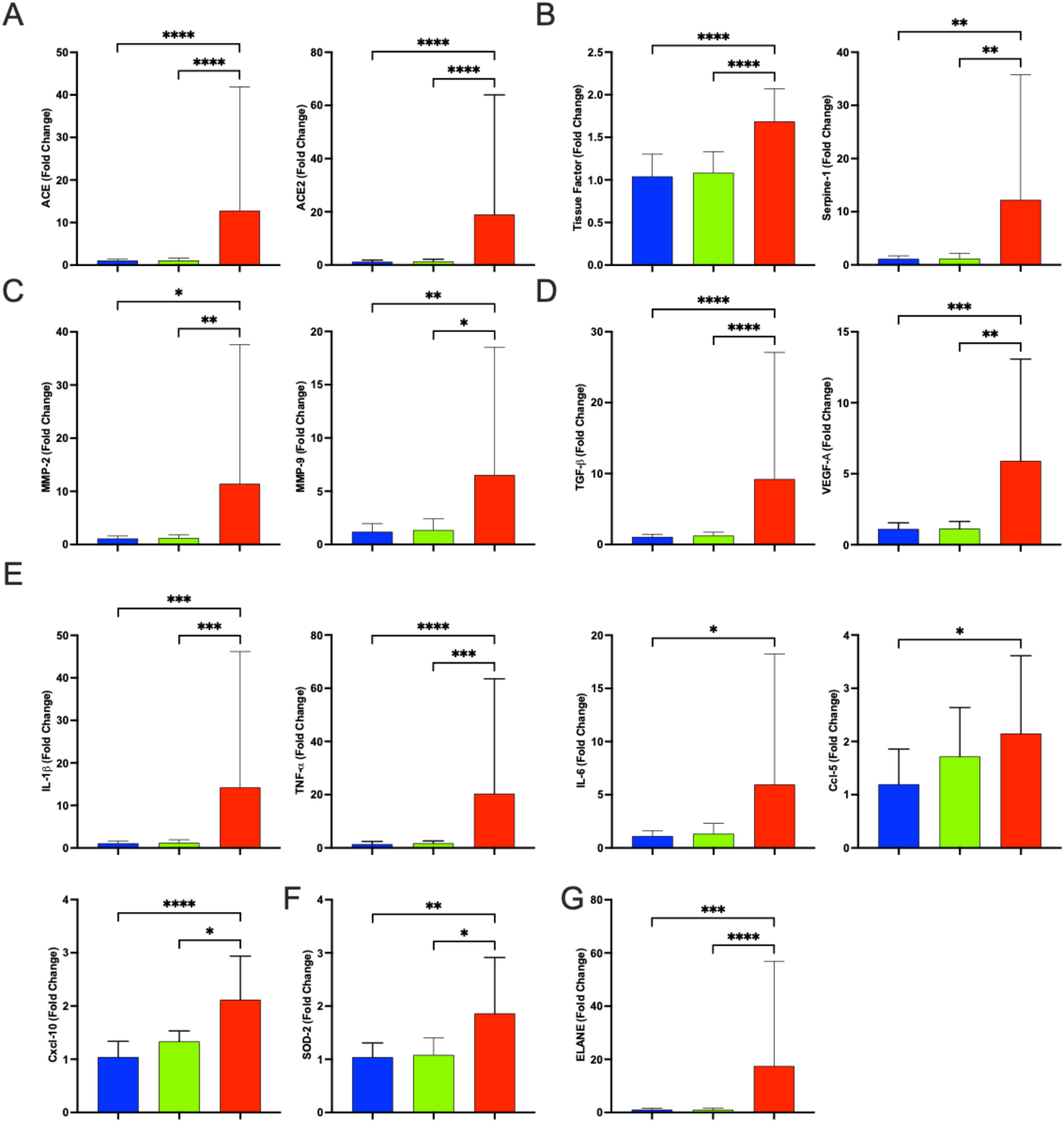
Acute e-cig exposure affects mRNA expression of target genes in the lung analyzed by RT-qPCR. Hamsters were exposed to bias flow (blue), PG/VG + 0% nicotine (green), and PG/VG + 2.5% nicotine (red) for two days (n= 14-16/ group). Target mRNA expression was analyzed by RT-qPCR. Expression was normalized to Rpl18 and calculated relative to the mean expression level of the biological replicates of the bias flow group. (a) RAAS pathway genes ACE and ACE2, (b) coagulation genes tissue factor and Serpine-1, (c) ECM remodeling genes MMP-2 and MMP-9, (d) fibrosis- and angiogenesis-/ vasculogenesis-related genes TGF-β and VEGF-A e) type 1 inflammatory genes IL-1β, TNF-β, IL-6, CCL-5, and CXCL-10, (f) ROS processing gene SOD-2, and (g) NETosis-related gene ELANE. *p<0.05, **p<0.01, ***p<0.001, ****p<0.0001, Kruskal-Wallis with Dunn’s post-hoc test.

## DISCUSSION

The purpose of this study was to determine if the golden Syrian hamster could be used to model the effects of e-cig vaping. In testing our hypothesis that hamsters are a viable model of e-cig induced airway inhalational injury, we exposed them to bias flow, PG/VG + 0% nicotine, and PG/VG + 2.5% nicotine. After exposure, we monitored exposure quality with serum cotinine, evaluated airway histopathology, and performed BALF studies. Additionally, we investigated the expression of 15 gene targets that covered pathways involving type 1 inflammation, processing of ROS, ECM remodeling, NETosis, fibrosis, and coagulation.

Our results fulfill the primary study objective that hamsters are a feasible model of the effects of acute e-cig vaping. First, we show that the measure of serum cotinine is an accurate and reproducible way to confirm exposure to nicotine vaping in hamsters (Fig. 1A). These levels are similar in magnitude to those reported for newborn mice exposed to 1.8% nicotine with PG for ten days and rats exposed to PG/VG + 2.5% nicotine for three days (7, 12). We evaluated histologic changes with acute vaping in the nose and lung. In the nose, AB-PAS staining suggested no change in mucin components of the epithelial layer (Fig. 1B). Although other studies have reported vaping-induced increases in MUC5AC, we suspect that our vaping protocol may not be long enough to induce changes visible on histological evaluation (13). In the lung, H&E staining demonstrated no change in bronchial epithelial organization or alteration in sloughing of cells into the airway lumen for vaped hamsters compared to bias flow controls. Finally, we evaluated BALF to examine the impact of acute vaping on total WBC counts and the percent of macrophages in that cell population. We did not see significant differences between our bias flow controls and vaping groups, with or without nicotine, which is consistent with a published report of acute vaping in a rat model (Fig. 1C) (7).

Most notably, the results indicated that there were increases in the nose and lung expression of genes of interest. Since hamsters are obligate nasal breathers and have a larger nasopharyngeal surface area compared to humans, nasopharyngeal studies were pursued. Our analysis of the nasal respiratory epithelium demonstrated nicotine-dependent increases in type 1 inflammation and fibrosis (Fig. 2A and B). There was a nicotine-independent increase in ROS processing (Fig. 2C). Additionally, there was a nicotine-independent decrease in the angiogenesis- and vasculogenesis-related gene VEGF-A (Fig. 2D); however, we are unsure of the clinical significance of this finding.

In the lung, the effects of vaping were highlighted by a nicotine-dependent increase in renin-angiotensin pathway by upregulation of ACE and ACE2 expression (Fig. 3A). The findings for ACE2 are similar to a published report which utilized a murine model of vaping (4). Importantly, we describe herein the novel finding of increased tissue factor expression due to nicotine vaping (Fig. 3B). Tissue factor activation is the initiator of the extrinsic coagulation cascade, and it has been implicated as a key to the process of microvascular thrombi formation and immunothrombosis in severe ARDS, including COVID-19 (14, 15). Although we have not elucidated the exact mechanism of the increase in tissue factor, our hypothesis is that it is due to the nicotine-dependent increases in type 1 inflammatory cytokine genes, which were also highly expressed in the lung tissue in our model (Fig. 3E). The expression of another coagulation-related gene, Serpine-1, was also markedly increased in the hamster lung following acute nicotine vaping (Fig. 3B). Serpine-1 encodes plasminogen activator inhibitor-1 (PAI-1), an inhibitor of fibrinolysis, which inhibits clot breakdown and pushes the coagulation system in a pro-coagulant direction (16).

Additional pathways demonstrating a nicotine-dependent response included NETosis, which was impacted by vaping through a nicotine-dependent increase in ELANE (neutrophil elastase) in the hamster lung (Fig. 3G). This gene encodes a serine protease whose expression is limited to neutrophils and their precursors. In cell culture, short term nicotine exposure led to increased ELANE gene expression in a myeloblast/ promyelocyte cell line (17). Subsequently, human BALF cells from individuals who chronically smoke or vape demonstrated increased neutrophil elastase expression (18). Since our acute exposure did not result in an influx of neutrophils into the lung, as measured by bronchoalveolar lavage and seen in histopathology, we suspect that this increase could be due to increased expression in the neutrophils already present in the tissue or from neutrophil precursors circulating in, and/or marginated within, the lung.

Our model also showed altered expression of genes involved in the development of lung fibrosis through nicotine-dependent increases in TGF-β and PAI-1 (Fig. 3 B and D). TGF-β is a principal regulator of lung fibrosis development, and its role in the development of lung fibrosis post-COVID infection is actively being studied. PAI-1 is also thought to have a role in lung fibrosis development through the induction of senescence in alveolar type II cells, beyond its role in fibrinolysis (19). Also potentially related to lung fibrosis, we report nicotine-dependent increases in MMP-2 and MMP-9, which are involved in ECM remodeling (Fig. 3C). Although other groups have shown a nicotine-independent decrease in MMP-2 and increase in MMP-9, the inconsistency may be due to species differences (4, 5). Overall, our findings implicate the potential for e-cig use to lead to dysregulated ECM repair. Finally, our findings in the lung demonstrate a differential response compared to the nose with a nicotine-dependent increase in ROS processing, angiogenesis, and vasculogenesis (Fig. 3 D and F).

In conclusion, the golden Syrian hamster has provided a viable model for the study of e-cig vaping-related airway injury and demonstrated significant upregulation of inflammatory pathways, even in an acute exposure. Through the development of this model, we will be able to further study potential sex differences in acute vaping exposure, in addition to the impact of e-cig use on SARS-CoV-2 infection. Its development allowed us to identify tissue factor as a novel nicotine-upregulated gene associated with vaping. This finding, along with the upregulation of other components of the type 1 inflammation and fibrotic pathways, are important targets of future vaping research.

## Supporting information

Supplemental Results

## ACKNOWLEDGEMENTS

This work was supported by the Countermeasures Against Chemical Threats (CounterACT) Program, National Institutes of Health (NIH), National Institute of Environmental Health Sciences (NIEHS) (U54ES027698-05 to CWW), and an associated Supplement (3U54ES027698-05S1). Additional support was provided by a grant from Children’s Hospital Colorado Research Foundation (CHC R7530S PEDS) (CWW). In addition, the authors are grateful for support of Dr. Robin R. Deterding, the Breathing Institute at Children’s Hospital Colorado, Dr. Livia A. Veress, and Jacqueline S. Rioux.

