## Supplemental Results for "Acute Vaping of a golden Syrian Hamster is Feasible and Leads to Nicotine-Dependent Respiratory Tract Inflammation"

**Supplement - Results**

**Supplemental Figure 1 (A-C)**

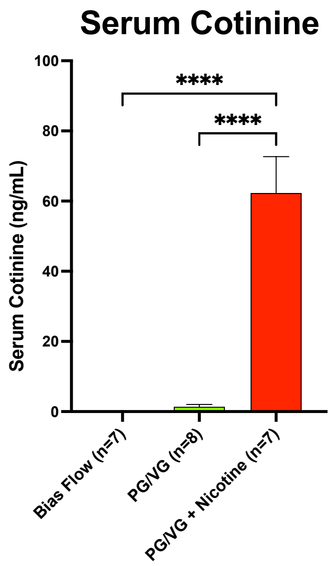


A

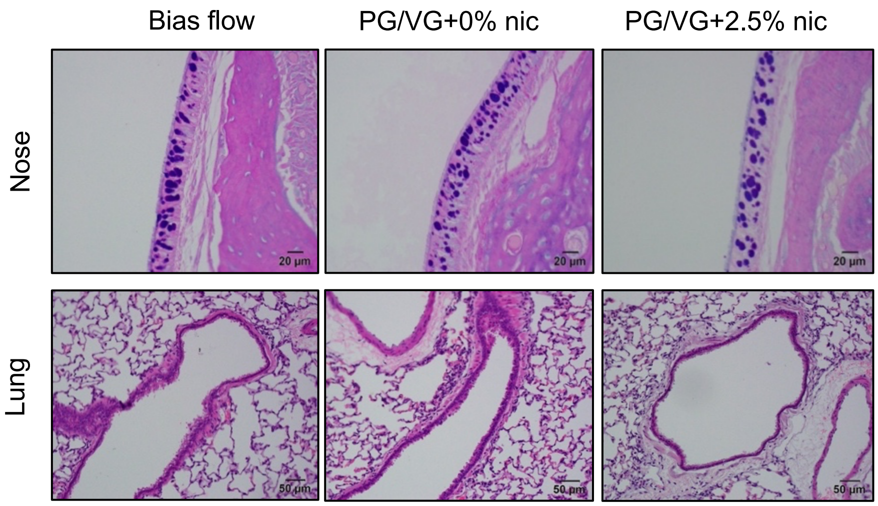


B

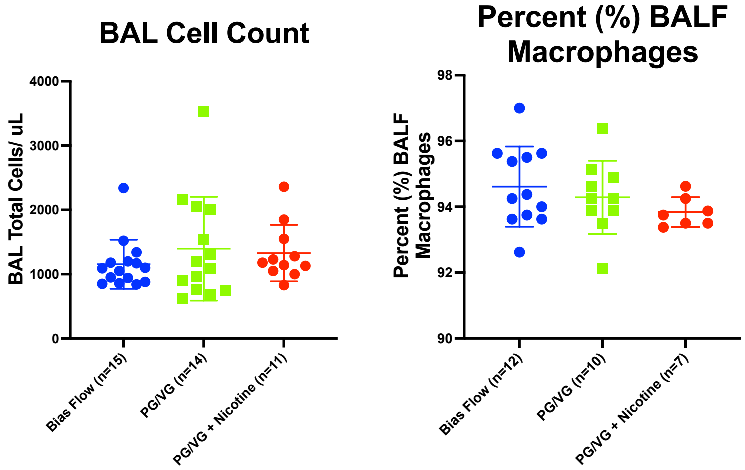


C

**Legend –Supplemental Figure 1** - Acute, whole-body e-cigarette exposure. a) serum cotinine levels for animals exposed to bias flow (0.0 ± 0.0 ng/mL), PG/VG + 0% nicotine (1.4 ± 0.7 ng/mL), or PG/VG + 2.5% nicotine (62.3 ± 10.4 ng/mL). **** $p < 0.0001$, ANOVA with Tukey-Kramer post-hoc test. b) AB-PAS-stained nasal sections and H&E-stained lung sections from animals exposed to bias flow (left panels), PG/VG + 0% nicotine (middle panels), or PG/VG + 2.5%

nicotine (right panels). c) total cell count and proportion of macrophages in bronchoalveolar lavage (BAL) fluid for animals exposed to bias flow (1154 ± 381 cells/uL), PG/VG + 0% nicotine (1396 ± 806 cells/uL), and PG/VG + 2.5% nicotine (1327 ± 439). BAL cell count p = 0.5171, percent BALF macrophages p = 0.3151, ANOVA with Tukey-Kramer post-hoc test. All data are reported as mean ± SD.

**Supplemental Figure 2 – Nose H&E; Lung AB-PAS**

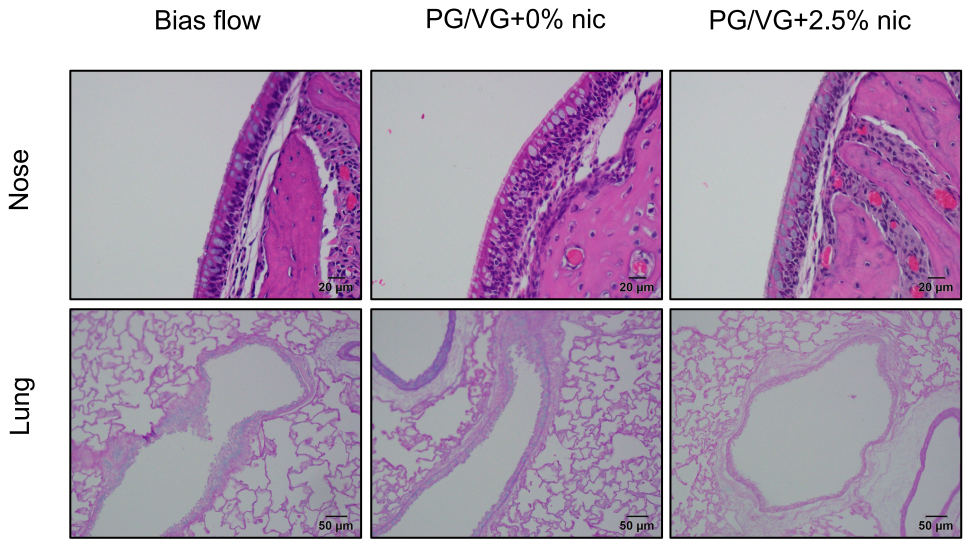


**Legend – Supplemental Figure 2 –** Hematoxylin and eosin (H&E) nasal sections and Alcian blue, Periodic acid-schiff (AB-PAS) lung sections from animals exposed to bias flow (left panels), PG/VG + 0% nicotine (middle panels), and PG/VG + 2.5% nicotine (right panels).

**Supplemental Figure 3**

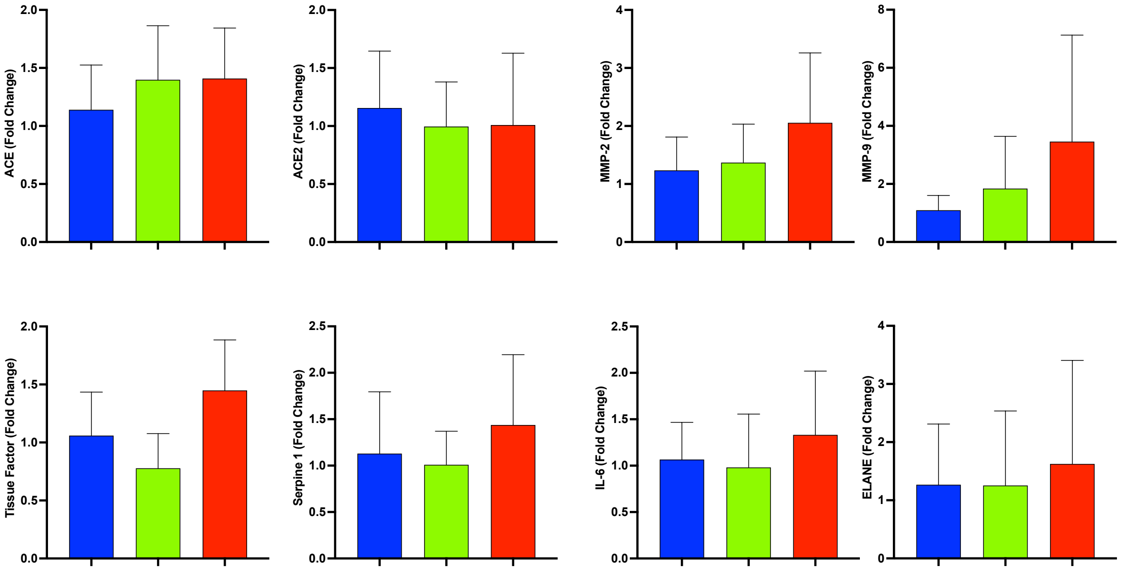


**Legend –Supplement Figure 3** - Acute e-cig exposure does not change mRNA expression of target genes in the nose analyzed by RT-qPCR. Hamsters were exposed to bias flow (blue), PG/VG + 0% nicotine (green), and PG/VG + 2.5% nicotine (red) for two days (n = 11-12/ group). Target mRNA expression was analyzed by RT-qPCR. Levels were normalized to Rpl18 and calculated relative to the mean expression level of the biological replicates of the bias flow group. There were no significant changes in ACE, ACE2, tissue factor, Serpine-1, MMP-2, MMP-9, IL-6, and ELANE. Kruskal-Wallis with Dunn's post-hoc test.
